# Drivers of HIV-1 transmission: the Portuguese case

**DOI:** 10.1101/655514

**Authors:** Andrea-Clemencia Pineda-Peña, Marta Pingarilho, Guangdi Li, Bram Vrancken, Pieter Libin, Perpetua Gomes, Ricardo Jorge Camacho, Kristof Theys, Ana Barroso Abecasis, on behalf of the Portuguese HIV-1 Resistance Study Group

## Abstract

**Background:** Portugal has one of the most severe HIV-1 epidemic in Western Europe. Two subtypes circulate in parallel since the beginning of the epidemic. Comparing their transmission patterns and its association with transmitted drug resistance (TDR) is important to pinpoint transmission hotspots and to develop evidence-based treatment guidelines.

**Methods:** 3599 HIV-1 naive patients collected between 2001 and 2014 were included in the study. Sequences obtained from drug resistance testing were used for subtyping, TDR determination and transmission clusters (TC) analyses.

**Results:** Subtype B transmission was significantly associated with young males, while subtype G was associated with older heterosexuals. Young males infected with subtype B were more likely to be included in TC. Consistently, a decreasing trend of prevalence and transmission of subtype G in Portuguese originated people was observed. Active TCs were associated with subtype B-infected males residing in Lisbon. TDR was significantly different when comparing subtypes B (10.8% [9.5-12.2]) and G (7.6% [6.4-9.0]) (p=0.001).

**Discussion:** TC analyses shows that the subtype B epidemic is active and fueled by young male patients residing in Lisbon and that transmission of subtype G in Portugal is decreasing. Despite similar treatment rates for both subtypes in Portugal, TDR is different between subtypes.

## 1. INTRODUCTION

Portugal had one of the highest rates of HIV diagnoses in Europe in 2016, with 10.0 diagnoses per 100,000 population [1]. Despite the fact that new diagnoses have decreased within the country in the last years [2], the patterns of HIV-1 transmission remain uncertain. Phylogenetic analyses are powerful tools to understand the dynamics of viral transmission [3–7], to provide insights for designing prevention policies.

According to the European and Portuguese guidelines for antiretroviral treatment [8,9], a baseline resistance test should be performed to determine transmitted drug resistance (TDR), which can impact the first-line antiretroviral response [10]. The last nationwide survey was carried out in 2003 and showed 7.8% of TDR [11]. Surveillance of TDR is important for the development of treatment guidelines, especially in Portugal where considerable migration from Portuguese speaking countries occurs, including some African countries where the levels of TDR are increasing along with the recent scaling-up of NRTI and NNRTI based treatments [12,13].

The epidemiology of HIV-1 in Portugal is unique in comparison to other European countries. Most of the epidemic is caused by parallel sub-epidemics of subtype B and subtype G [14]. Until 2005, subtype B accounted for approximately 40% of infections and subtype G accounted for 30% [11,14,15]. The present large-scale cohort provides the unique opportunity to compare the temporal evolution of the parallel epidemics of these subtypes in the same country. Herein, we use transmission cluster reconstruction to understand the drivers of HIV-1 transmission in Portugal and its correlation with primary drug resistance: prevalence of TDR and factors associated with the spread of TDR. The characterization of HIV-1 transmission in the Portuguese epidemic can help to design targeted prevention strategies.

## 2. PATIENTS AND METHODS

### Study Population

The protocol was in accordance with the Declaration of Helsinki and approved by Ethical Committees of Centro Hospitalar de Lisboa Ocidental (108/CES-2014). The Portuguese HIV-1 drug resistance database contains clinical and genotype resistance testing data from patients followed up in 22 hospitals located around the country [16]. The inclusion criteria for the analysis of TDR was age older than 18 years and no history of antiretroviral treatment between January 2001 and December 2014. This cohort is named PT-naive, hereafter. The genomic data included the protease and the reverse transcriptase (HXB2: 2253-3554) obtained through population sequencing using the ViroSeq assay.

### Drug Resistance Assessment

Surveillance drug resistance mutations (SDRM) were defined according to the WHO list [17]. The impact of TDR was evaluated with the HIVdb v.7.0 and Rega v.9.1.0 (http://sierra2.stanford.edu/sierra/servlet/JSierra?action=algSequenceInput).

### Subtyping and Transmission Cluster Analyses

HIV-1 subtypes were determined with Rega v3 and COMET v.1.0 [18] [19,20]. Subtype G and CRF14_BG were merged in a single group (named hereafter G dataset), given that: i) this genomic region has the same evolutionary origin for G and CRF14_BG strains; ii) the origin of the CRF14_BG strains occurred in the Iberian Peninsula [14,19,21]; iii) previously, we reported that the two tools and the manual phylogenetic analyses were not conclusive whether the sequences were G or CRF14_BG in the present cohort [17]; iv) there was no recombination breakpoint in the genomic regions of protease and reverse transcriptase in these sequences. A statistical sub-analysis was performed considering only “pure" subtype G strains (excluding CRF14_BG), defined by the concordant assignment of the two subtyping tools [19,20], to evaluate how this affected our findings (Supplementary material).

For the TCs analysis, the dataset was complemented with controls retrieved from: (i) the treated population of the Portuguese cohort between 2001 and 2014, (ii) the 50 best-matched sequences to each sequence of the total cohort of subtype B and G, as retrieved by BLAST (http://blast.ncbi.nlm.nih.gov/Blast.cgi), (iii) all HIV-1 *pol* subtype B and G sequences available from Portugal in the Los Alamos database (http://www.hiv.lanl.gov) [22]. Three subtype D or B reference sequences were used as the outgroup. Sequences with low quality, duplicates and clones were deleted. The resulting dataset was aligned with Muscle [23] and verified for codon-correctness using VIRULIGN [24]. To avoid convergent evolution, SDRMs were removed [17]. The final subtype B and G datasets consisted of 7497 and 4372 sequences, both with a length of 1173 nucleotides (IQR:1173-1173).

A Maximum likelihood tree was constructed with the GTR+ 4Γ nucleotide substitution model and 1000 bootstraps, as implemented in RAxML version 7.5.5. ^21^ The transmission clusters (TC) were identified with Cluster Picker using a threshold that included a genetic distance of 0.045 and ≥ 80% bootstrap replicates [5,25]. To evaluate the effect of the definition of TCs in the results, sensitivity analyses were performed with varying genetic distances (0.015, 0.030, 0.045, 0.060) and bootstrap supports (70, 90, 95, 98).

TCs identified were confirmed with Bayesian Markov Chain Monte Carlo (MCMC) inference, as implemented in BEAST v1.8.2 [26]. The temporal signal of the TCs datasets was evaluated with TempEst [27]. The uncorrelated log-normal relaxed molecular clock with a discretized GTR substitution model and the Bayesian Skygrid coalescent model were used [28]. Three separate MCMC chains were run for at least 100 million generations. Convergence was determined with Tracer using a burn-in of 10% (http://beast.bio.ed.ac.uk/Tracer). The maximum clade credibility (MCC) tree was constructed with TreeAnnotator after discarding the burn-in, and visualized with FigTree v1.4.2 (http://tree.bio.ed.ac.uk).

The TCs analyses included the following definitions: (i) A pair was defined as exactly two patients included in a TC, one of them from the PT-naive cohort; (ii) a cluster ≥3 included three or more patients, with at least one from the PT-naive cohort (iii) a TDR-cluster-≥3 or TDR-pair contain at least one PT-naive patient with a sequence harboring a SDRM; (iv) an onward-TDR-cluster had ≥3 patients with the same SDRM in the majority of the patients and at least one from the PT-naive cohort with TDR, which suggest onward transmission of TDR; and (v) active-TCs included transmission of HIV-1 or/and TDR that involves at least one PT-naive patient within a time frame of ≤5 years. The time frame was calculated as the maximum length of time between the ancestral node and the most recent tip (year 2014) of the MCC trees [29]. As such, a TC could be separated in two or more active sub-clusters, since such sub-clusters may indicate the population which actively transmitted HIV-1 or TDR in the last years.

### Statistical analyses

Statistical analyses were performed to understand and compare the dynamics of the subtype B and G sub-epidemics in the PT-naive cohort, specifically the factors associated with transmission of HIV-1B and HIV-1G, independently of TDR; and the factors associated with transmission of TDR. All these analyses were performed within and compared between subtypes. Sensitivity analyses were performed and, if a result was discordant in the sensitivity analysis, the difference is clearly stated throughout the manuscript.

The Fisher’s exact test or regression techniques were conducted to compare between proportions, while the Mann Whitney U test or the t-test were used to compare between median or mean values for continuous variables, as appropriate. Binomial logistic regression was used to determine the factors associated with each epidemic, TDR, and clustering. The Bonferroni method was used for multiple testing adjustments. The level of statistical significance was set at 5%. The analyses were performed with the statistical R software v.3.2.1.

## 3. RESULTS

### 3.1. Study Population

The PT-naive cohort included a total of 3599 patients, 2042 with subtype B (56.7%) and 1557 (43.3%) with G. The socio-demographic factors are shown in Supplementary Tables 1&2.

### 3.2. Subtype B sub-epidemic is associated with young males living in Lisbon

Regarding the socio-demographic factors associated with the transmission of the sub-epidemic B versus G in the PT-naive cohort, there were significant differences between the two sub-epidemics for age, gender, risk of transmission, residence in Lisbon, and CD4 count in the univariate analyses (Supplementary Tables-1&2), while in the multivariate analyses younger age (Odds Ratio (OR): 0.83 for every increase of 10 years (^10-years^), 95% Confidence interval: 0.79-0.89, p<0.0001), male (OR: 2.66, 2.28-3.09, p<0.0001) and living in Lisbon (OR: 1.44, 1.25-1.66, p<0.0001) were significantly associated with subtype B infections

### 3.3. Transmission of subtype B is driven by young males

There were 497 TCs that included 61.2% of the subtype B PT-naive cohort (Table 1, and Supplementary Tables 1&2). When comparing the cohort outside versus inside TCs of subtype B in the multivariate analysis, individuals inside subtype B TCs were younger (OR^10-years^: 0.83, 0.76-0.90, p<0.0001) and more frequently male (OR 1.42, 1.14-1.78, p=0.001) than individuals outside the TCs, indicating that young males are driving this sub-epidemic in Portugal as we have identified in other European cohorts[5].

**Table 1:**
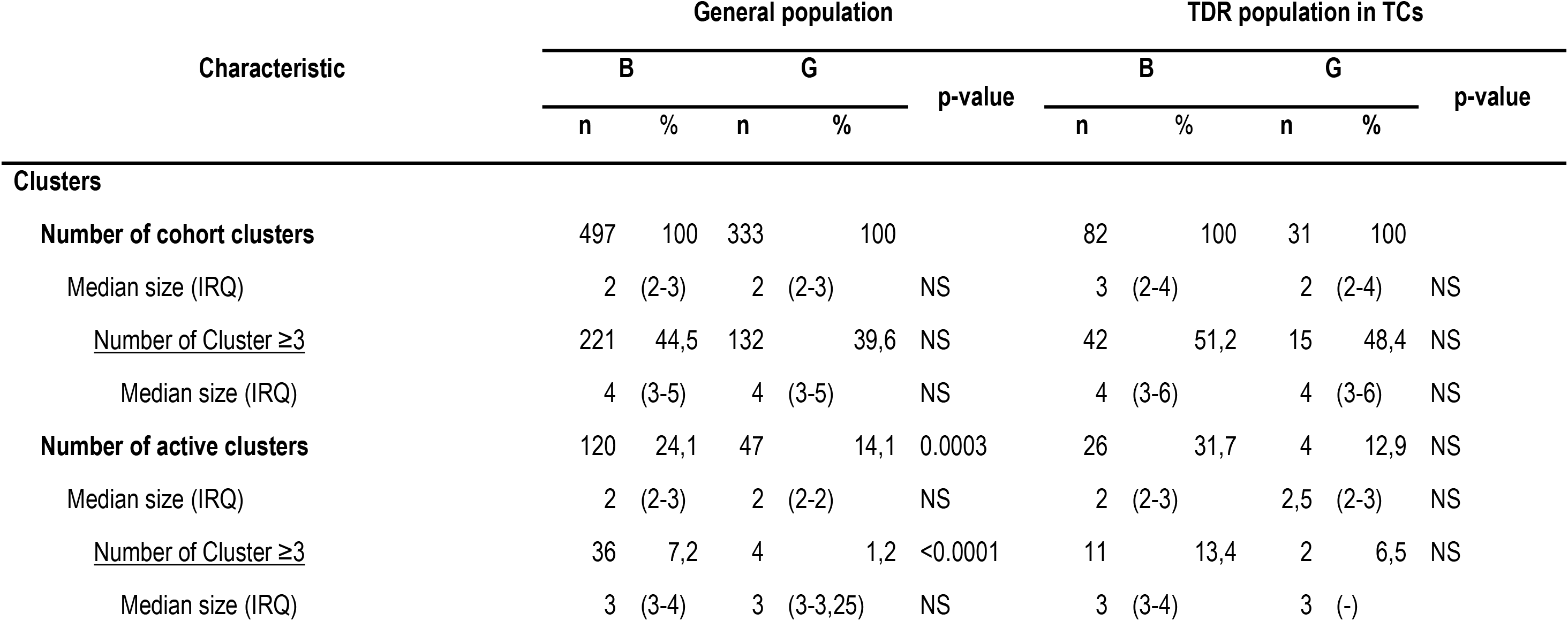
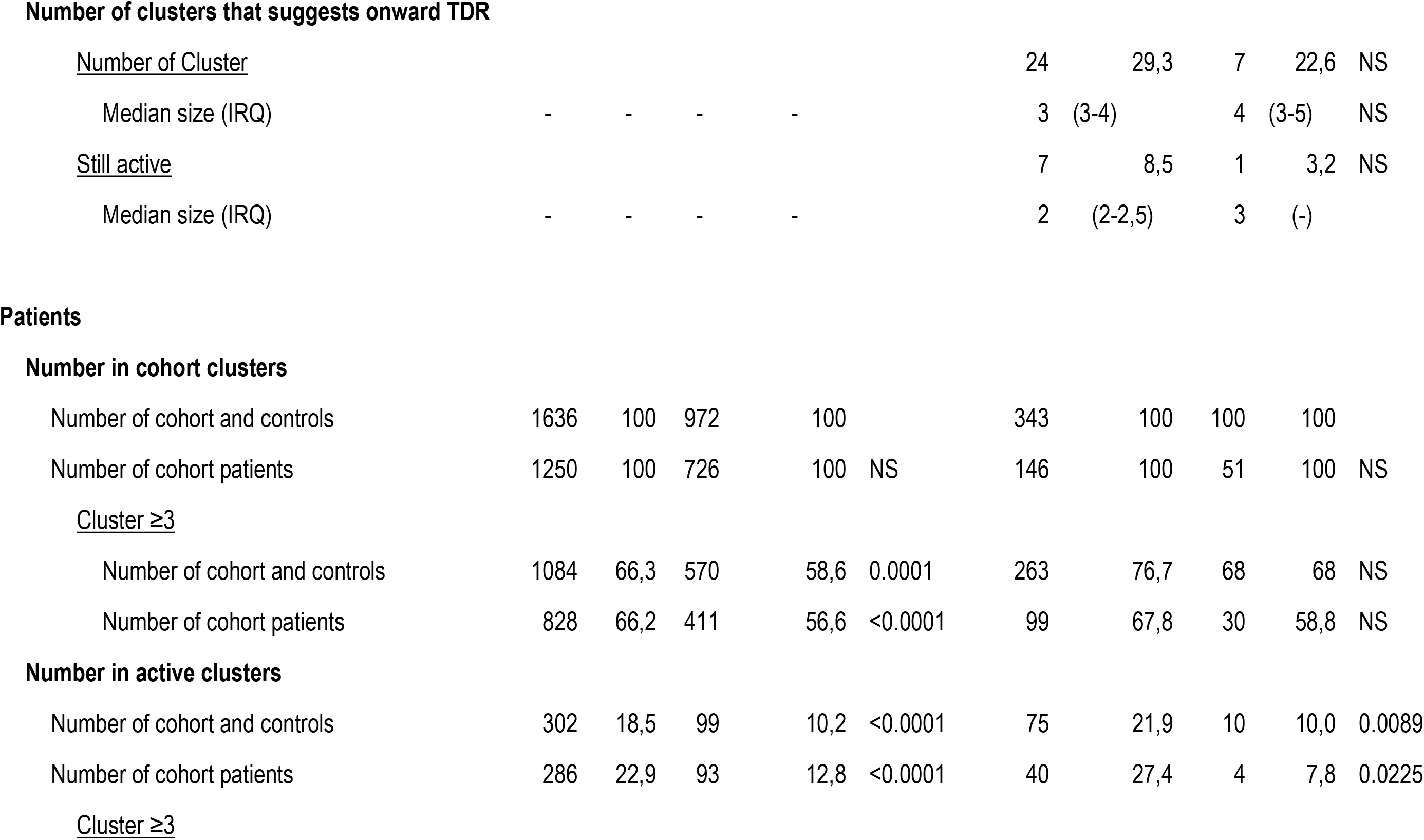
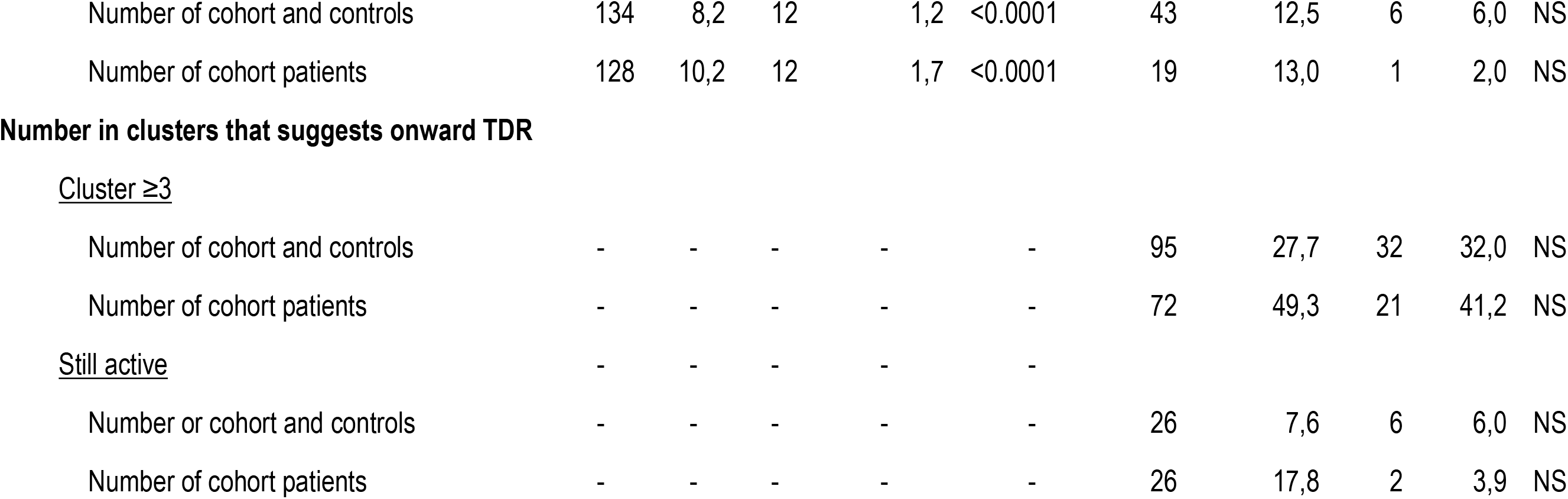
Characteristics of the transmission clusters (TCs) in the general cohort and in the cohort with TDR. Abbreviations: IQR: interquartile range, n: sample, NS: No significant, TCs: Transmission clusters TDR: transmitted drug resistance, % percentage.

The number of PT-naive patients included in TCs had a peak in 1999 for clusters ≥3 followed by a steady decrease, while the peak in the number of patients in pairs occurred in 2002 followed by an up and down curve (Figure-1A). This peak in 1999 includes 38% of the PT-naive cohort, which suggests transmission of HIV was still ongoing for subtype B despite the introduction of HAART.

**Figure 1:**
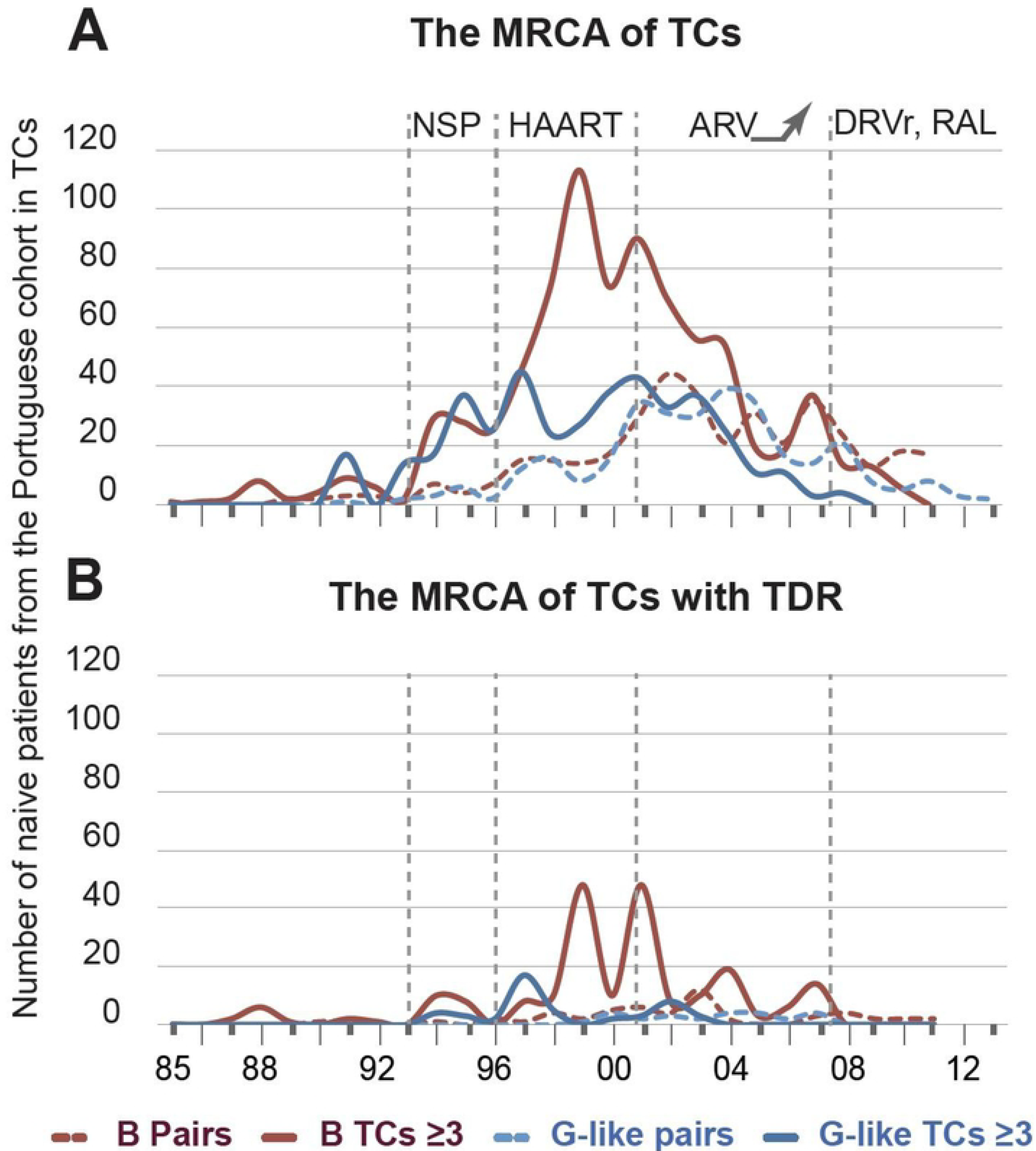
Prevalence in percentage of G subtype in Portuguese originated people during the period of the study (years in the y-axis) when considering the ones who were in transmission clusters (**A**). The light blue shades are the confidence intervals of the proportion of patients for a period of two years (dark blue line). The first period was excluded given the few number of patients but significance did not change. (**B**) Prevalence (dot) and 95% confidence intervals of the Transmitted drug resistance for subtypes B (red) and G (dark blue) for the PT-naive cohort and for each drug group. Geographical differences were observed for TDR when Lisbon was compared with other regions. Significant differences are shown with an asterisk. Abbreviations: NRTI: Nucleoside reverse transcriptase inhibitors, NNRTI: non-NRTI, PI: Protease inhibitors; TCs: Transmission clusters, vs: versus, %: percentage.

### 3.4. Transmission of subtype G is decreasing in native Portuguese people

There were 333 TCs that contained 46.6% of the G PT-naive cohort (Table 1, and Supplementary tables 1&2). None of the socio-demographic or clinical factors were significantly associated with transmission of subtype G. Interestingly, a decreasing trend in the percentage of native Portuguese people included in TCs was observed since 2005 (p=0.006, Figure-2A).

**Figure 2:**
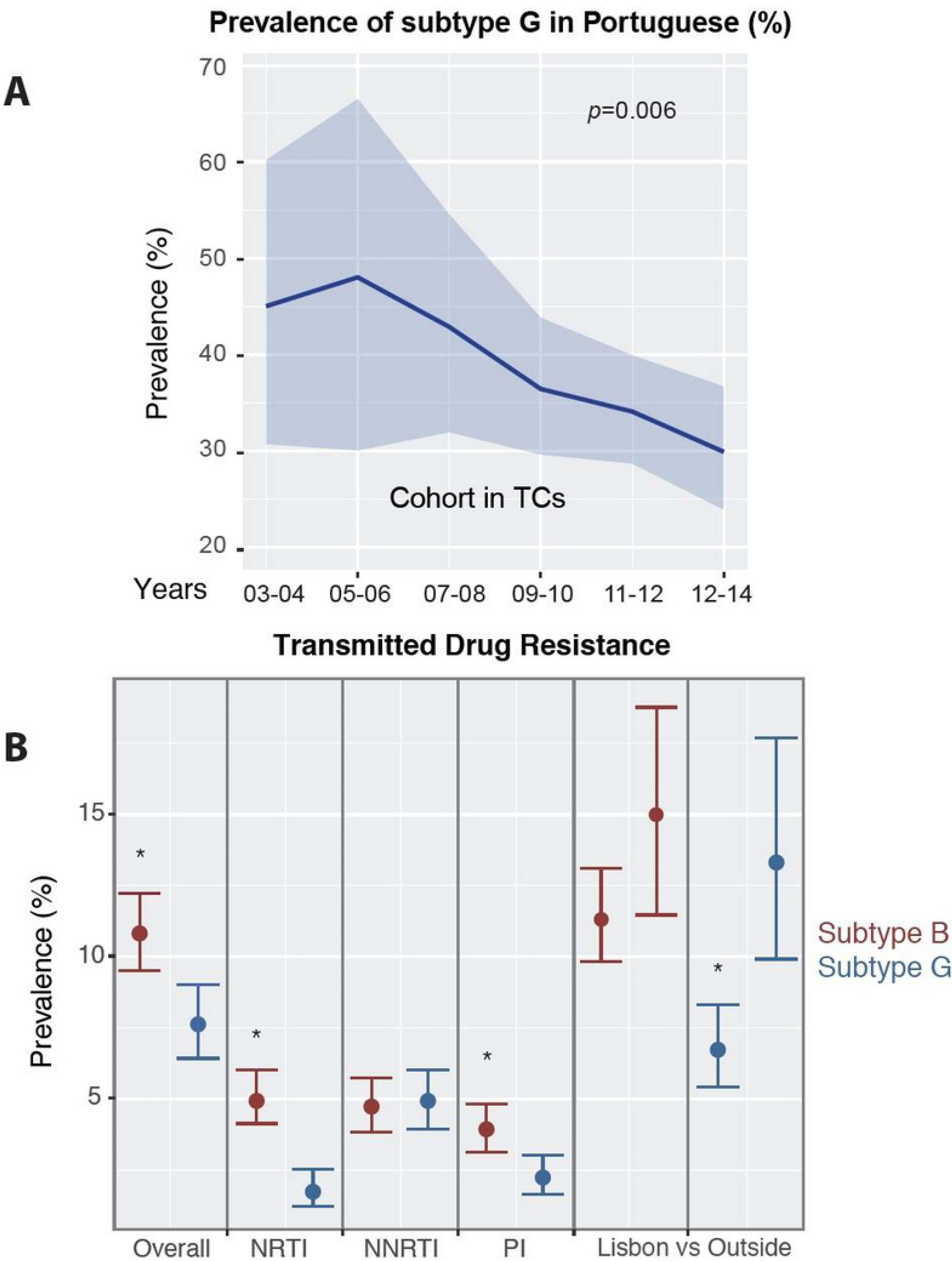
The most recent common ancestor (MRCA) or the origin of the transmission clusters in years (y-axis) in the PT-naive cohort. The number of naive patients included in the TCs is shown in the y-axis while the year of origin is in the x-axis. The type of clusters is presented as pairs (dashed line) and clusters ≥3 (solid line) for B (dark red) and G-like (blue). (**A**) The number of patients from the PT-naive cohort is shown according to the MRCA of the TCs (**B**) The number of patients from the PT-naive cohort is shown according to the MRCA of the TCs with TDR. The Highly Effective Antiretroviral Therapy (HAART) was introduced in 1996. The needle and syringe program (NSP) was introduced in Portugal in 1993. There were several antiretroviral (ARV) drugs introduced in the earlier 2000s such as Tenofovir, Emtricitabine and Protease inhibitors such as Lopinavir/ritonavir, Fosamprenavir/ritonavir and Atazanavir/ritonavir, which increased the regimen options. Since 2007 onwards, potent drugs such as Darunavir (DRV) and the first integrase inhibitors - Raltegravir (RAL) were introduced.

The numbers of PT-naive patients included in TCs had an up and down curve, as clusters ≥ 3 originated more frequently in 2001 followed by a steady decrease, while this peak occurred in 2005 for pairs (Figure-1A). In contrast with subtype B, most PT-naive patients were involved in TCs originated before the introduction of HAART; i.e. G: 45 TCs included 266 patients (17.1%) vs B: 63 TCs with 114 patients (9%);(p<0.0001).

### 3.5. In the last years, subtype B transmission was predominant and occurred between patients sampled in Portugal

In a sub-analysis considering only TCs originating in the last five years of the cohort (active-TCs), 24.1% (120/497) and 14.1% (47/333) TCs were subdivided in smaller active-TCs for subtype B and G, respectively (Table-1). These active-TCs included mainly subtype B patients (75,5%, 286/379 compared to 23.9% (93/379) for subtype G (p<0.0001)). Socio-demographic characteristics of the patients in active-TCs and the total cohort were similar (Supplementary Tables 1&2). Males (OR: 6.56, 3.63-11.85, p<0.0001) and patients living in Lisbon (OR: 2.03, 1.12-3.69, p<0.05) were associated with the active transmission of subtype B when compared with G. Since we completed our cohort with controls retrieved from other databases, it is important to note the active-TCs included mainly controls sampled in Portugal for both subtypes, indicating transmission of HIV-1 predominantly occurs between patients in Portugal.

### 3.6. TDR in subtype G occurs more frequently in patients followed-up in hospitals outside the Lisbon area

In the subtype B PT-naive cohort, the prevalence of TDR was 10.8% [221/2042; 9.5-12.2%], 4.9% [102/2042; 4.1-6.0] for nucleoside reverse transcriptase inhibitors (NRTIs), 4.7% [96/2042; 3.8-5.7%] for Non-NRTIs (NNRTIs) and 3.9% [80/2042; 3.1-4.8] for protease inhibitors (PIs) (Figure-2B). Regarding the socio-economical and clinical factors, none of the factors were associated with TDR in the multivariate analysis for this subtype.

For subtype G, the TDR prevalence in the PT-naive cohort was 7.6% [118/1557, 6.4-9.0], 1.7% [27/1557; 1.2-2.5] for NRTIs, 4.9% [76/1557; 3.9-6.0] for NNRTIs and 2.2% [34/1557; 1.6-3.0] for PIs. Older age, heterosexual transmission and living outside of *Lisboa* and *Vale do Tejo* regions were significantly associated with TDR in subtype G (Figure-2B). The multivariate analysis showed people living outside of the *Lisboa* and *Vale do Tejo* regions associated with TDR in subtype G (OR: 1.87, 1.22-2.88).

When subtype B and G were compared, there were higher prevalence of TDR (OR: 1.47, 1.17-1.89, p=0.001), NRTIs TDR (OR: 2.98, 1.92-4.76, p <0.0001) and PIs TDR (OR: 1.83, 1.20-3.83, p=0.003) for subtype B. Subtype B patients with TDR were also older (OR^10-years^: 0.68, 0.55-0.82, p= 0.001) and more frequently male (OR: 3.10, 1.83-5.24, p<0.0001).

### 3.7. Active and onward transmission of TDR for subtype B is driven by males living in Lisbon

Eighty-two subtype B TCs had at least one patient from the PT-naive cohort harboring viruses with SDRMs (TDR-TCs, Table 1). The TDR-TCs included 66.1% (n=146/221) of the total number of patients with TDR compared to 43.2% for G (n=51/118; p<0.0001; Supplementary Table-1), indicating more active transmission of TDR in subtype B. When the Portuguese treated population was included as complementary database for the TCs analyses, it was observed that nearly half (n=39/82) of the TDR-TCs for subtype B included at least one treated patient. However, the number of treated patients in subtype B TDR-TCs decreased over time since 2006 (p<0.0001).

The origin of subtype B TDR-TCs was mainly between 1999 and 2005: 23 pairs and 26 clusters ≥3 represented 60% (n=88/146) of the PT-naive cohort in clusters harbouring viruses with SDRMs (Figure-1B). There were no socio-demographic factors associated with the transmission of SDRMs for subtype B.

Twenty-six TDR-TCs originated in the last five years for subtype B (active clusters; Table-1 and supplementary Table-1). This population was similar to the population involved in subtype B active clusters: male (85%, 64/75), living in *Lisboa* and *Vale do Tejo* region and <35 years old (both 65.3%). Twenty-four TDR-TCs had evidence of onward transmission of SDRMs, those were mainly thymidine analog mutations (TAMs) and/or NNRTIs SDRMs. Seven out of those 24 TDR-TCs with evidence of onward transmission were still active in the last five years. The characteristics of the population reflected that male (77%, 20/26) and living *Lisboa* and *Vale do Tejo* region (69.2%) still drive the transmission of TDR.

As expected, when the two subtype B and G sub-epidemics transmitting drug resistance were compared, subtype B was significantly associated with TDR transmission (OR: 1.75, 1.24-2.49, p=0.0007). Younger age (OR^10-years^: 0.53, 0.39-0.73, p<0.0001) and males (OR: 4.53, 1.92-10.66 p=0.0005) were consistently associated with transmission of subtype B TDR in all analyses.

### 3.8. The onward and active transmission of TDR for subtype G is limited

Thirty-one subtype G TDR-TCs included 43.2% (51/118) patients from the PT-naive cohort harbouring viruses with SDRMs (Table 1 and Supplementary Table-2). Nearly half (n=15/31) of the TDR-TCs for subtype G included at least one treated patient.

When considering the time origin of TDR-TCs, 35.3% patients of the PT-naive cohort were involved in TDR-TCs originated between 1996 and 1999, followed by 27.5% between 2000-2003 (Figure-1B). Then, mainly pairs including 27.5% and 25.5% of the TDR-patients were originated in 2000-2003 and 2004-2008, respectively. There were no socio-demographic factors or time trends associated with transmission of TDR. When including controls and comparing people transmitting TDR versus without TDR in TCs of subtype G, age (median: 44, IQR: 34-54 versus 37, 31-46, p=0.002), and viral load were significant in the univariate analysis (median: 4.9 Log-copies/mm^3^, IQR: 4.4-5.7 versus 4.6, 3.9-5.2, p=0.03). However, those were no longer significant in the multivariate analyses. When TDR-TCs ≥3 were analysed including controls, residence outside of the *Lisboa* and *Vale do Tejo* region was significantly associated with TDR within subtype G in the multivariate analysis (OR: 2.96, 1.29-6.79, p=0.01). Interestingly, this geographical pattern was no longer observed in the seven onward-TDR clusters, and from those only one was an active-TC. Unlike subtype B, the socio-demographic factors did not show any clear pattern.

## 4. DISCUSSION

To our knowledge this is the first study that uses phylodynamics to describe the drivers of HIV-1 transmission in Portugal. Transmission cluster reconstruction has been previously used to understand HIV-1 and resistance transmission patterns in other settings [3–7]. Herein, we combine and compare the information provided by classical statistical analyses of the most complete Portuguese cohort available, stratified by subtypes, with the one retrieved from transmission clusters analyses.

Through our detailed analyses of the Portuguese HIV-1 epidemic, we find strong indications that: 1) transmission of subtype B is associated with younger males; 2) transmission of subtype G is decreasing in Portugal and in the native Portuguese population; 3) transmission of drug resistance has different patterns: males living in *Lisboa* and *Vale do Tejo* regions drive the active and onward transmission of TDR, while this transmission is limited for subtype G and does not correlate with any socio-demographic factors.

Importantly, active transmission of subtype B in the last years has been driven by males residing in Lisbon. Although a source of uncertainty is the lack of risk factor information for a large part of our cohort, the consistency of our findings in different analyses suggest an important role of MSM living in Lisbon for this sub-epidemic. These results are consistent with other studies in Europe, Brazil or USA, where young MSMs have been identified as the main drivers of subtype B and TDR transmission [3–5,7]. More studies are needed to evaluate how tourism or migration may influence these results.

We observed a decline both in the prevalence and in the number of patients present in subtype G TCs in native Portuguese people since 2005 (Figure-2A). This indicates that transmission of subtype G strains is decreasing and that transmission has been limited in the last years. This sub-epidemic was unique in Europe and mainly circulating within Portugal. It was imported from West Africa, it was associated with intravenous drug users (IDUs) [30] and afterwards became also prevalent in heterosexuals. With the introduction of the needle and syringe program in 1993, new infections in IDUs declined and this could be potentially associated with the decrease of viral transmission together with the introduction of HAART in 1996 [2,31]. This finding corroborates how a long-term effective prevention program impacted HIV-1 transmission.

While we described that TDR levels differ between subtypes, with higher levels for B, the overall TDR remains stable across time, which agrees with the European study SPREAD [32]. TC analyses indicates that onward transmission of TDR is limited and mainly associated with subtype B and with a decreasing proportion of involvement of treated population since 2006. A higher TDR level for subtype B for NRTIs and PIs could result from several factors: i) The earlier beginning of the treatment for subtype B patients than for subtype G, and/or ii) a lower fitness of G strains in presence of SDRMs, which would cause faster reversion and consequent lower transmissibility level of SDRMs; and/or ii) behavioral patterns affecting the G sub-epidemic dynamics, with slower transmission rates and therefore higher likelihood that SDRM revert before their onward transmission; and/or iv) different treatment strategies for each subtype, which is unlikely because these patients are treated in the same country with similar regimens.

Our results should be interpreted with caution due to the lack of information about the time of infection, risk of transmission, country of origin and limited representativeness for the North region of Portugal [2]. The BEST-HOPE project is prospectively collecting recent socio-demographic and behavioral data to complete the picture of the current patterns of transmission in the country [33]. Finally, phylogenetic analyses have intrinsic limitations since it does not provide information about sexual networks and depends on the sampling density[34].

In conclusion, we have shown different patterns of transmission of HIV-1 and resistance for the two most important sub-epidemics in Portugal: subtype B and G. Our findings suggest that long-term prevention policies have impacted the transmission of subtype G in Portugal and resulted in decrease of prevalence of this subtype, while subtype B is reflecting the current patterns of HIV-1 transmission that is happening in other European countries.

## AUTHORS’ CONTRIBUTIONS

Conceptualization, Andrea Pineda-Peña and Ana Abecasis; Data curation, Andrea Pineda-Peña and Marta Pingarilho; Formal analysis, Andrea Pineda-Peña, Marta Pingarilho, Guangdi Li, Bram Vrancken, Pieter Libin, Kristof Theys and Ana Abecasis; Funding acquisition, Ana Abecasis; Investigation, Andrea Pineda-Peña, Perpetua Gomes, Ricardo Camacho and on behalf of the Portuguese HIV-1 Resistance Study Group; Methodology, Andrea Pineda-Peña, Guangdi Li, Bram Vrancken and Ana Abecasis; Project administration, Ana Abecasis; Resources, Perpetua Gomes, Ricardo Camacho and on behalf of the Portuguese HIV-1 Resistance Study Group; Supervision, Ana Abecasis; Validation, Marta Pingarilho; Writing – original draft, Andrea Pineda-Peña and Ana Abecasis; Writing – review & editing, Andrea Pineda-Peña, Marta Pingarilho, Pieter Libin, Kristof Theys and Ana Abecasis.

## ACKNOWLEDGEMENTS

We would like to thank the patients and all the members of the Portuguese HIV-1 Resistance Study Group:

Fátima Gonçalves, Isabel Diogo, Joaquim Cabanas, Ana Patrícia Carvalho, Sandra Fernandes, Inês Costa, Kamal Mansinho, Ana Cláudia Miranda, Isabel Aldir, Fernando Ventura, Jaime Nina, Fernando Borges, Emília Valadas, Manuela Doroana, Francisco Antunes, Maria João Aleixo, Maria João Águas, Júlio Botas, Teresa Branco, José Vera, Inês Vaz Pinto, José Poças, Joana Sá, Luís Duque, António Diniz, Ana Mineiro, Flora Gomes, Carlos Santos, Domitília Faria, Paula Fonseca, Paula Proença, Luís Tavares, Cristina Guerreiro, Jorge Narciso, Telo Faria, Eugénio Teófilo, Sofia Pinheiro, Isabel Germano, Umbelina Caixas, Nancy Faria, Ana Paula Reis, Margarida Bentes Jesus, Graça Amaro, Fausto Roxo, Ricardo Abreu and Isabel Neves.

## AVAILABILITY OF DATA

Access to the sequences linked to all clinical and demographic data is available through Euresist (http://www.euresist.org), a project aiming to collect and make available data to study HIV drug resistance and viral diversity in Europe.

